# Tissue-specific *trans* regulation of the mouse epigenome

**DOI:** 10.1101/322081

**Authors:** Christopher L. Baker, Michael Walker, Seda Arat, Guruprasad Ananda, Pavlina Petkova, Natalie Powers, Hui Tian, Catrina Spruce, Bo Ji, Dylan Rausch, Kwangbom Choi, Petko M. Petkov, Gregory W. Carter, Kenneth Paigen

## Abstract

Although a variety of writers, readers, and erasers of epigenetic modifications are known, we have little information about the underlying regulatory systems controlling the establishment and maintenance of the epigenetic landscape, which varies greatly among cell types. Here, we have explored how natural genetic variation impacts the epigenome in mice. Studying levels of H3K4me3, a histone modification at sites such as promoters, enhancers, and recombination hotspots, we found tissue-specific *trans*-regulation of H3K4me3 levels in four highly diverse cell types: male germ cells, embryonic stem (ES) cells, hepatocytes and cardiomyocytes. To identify the genetic loci involved, we measured H3K4me3 levels in male germ cells in a mapping population of 60 BXD recombinant inbred lines, identifying extensive *trans*-regulation primarily controlled by six major histone quantitative trait loci (hQTL). These chromatin regulatory loci act dominantly to suppress H3K4me3, which at hotspots reduces the likelihood of subsequent DNA double-strand breaks. QTL locations do not correspond with enzyme known to metabolize chromatin features. Instead their locations match clusters of zinc finger genes, making these possible candidates that explain the dominant suppression of H3K4me3. Collectively, these data describe an extensive, tissue-specific set of chromatin regulatory loci that control functionally related chromatin sites.

## INTRODUCTION

Both cellular differentiation and the functional capacities of differentiated cells are controlled by epigenetic DNA and histone modifications associated with a variety of genomic regulatory elements, including promoters, enhancers and insulators. Although much is known about the proteins that act as writers, readers, and erasers of these marks (reviewed in (Allis et al. 2015)), we know less about the underlying regulatory systems controlling their sites of action and tissue specificity. Understanding the properties of these regulatory systems is a matter of considerable significance as the majority of the known genetic variation affecting human health and disease is regulatory in nature (Maurano et al. 2012; Pickrell 2014) and chromatin changes are major features of both tumorigenesis (Verma et al. 2014; Avgustinova and Benitah 2016) and aging (Fraga et al. 2005; Martin 2005; Maegawa et al. 2014).

One avenue for exploring these questions is analyzing the genetic basis of natural variation in the epigenetic landscape (Albert and Kruglyak 2015; Lappalainen 2015; Taudt et al. 2016). Such variation can arise from mutations that act locally by altering the binding sites of transcription factors, which in turn influence levels of DNA or histone modifications (Kasowski et al. 2013; Kilpinen et al. 2013; McVicker et al. 2013). Local effects are typically *cis*-acting; the genetic determinant maps close to the location of the molecular phenotype it controls and only influences its own chromosome when heterozygous. The limitations of genetic mapping using human cell culture systems have largely restricted genetic studies of chromatin regulation to local *cis* effects (Albert and Kruglyak 2015; Pai et al. 2015; Taudt et al. 2016).

Alternatively, diffusible factors encoded by one region of the genome can act at a distance, regulating epigenetic modifications and gene expression at multiple sites in *trans*. By examining several chromatin marks in tissue from 30 rat recombinant inbred lines, one recent study found tissue-specific *trans*-regulation of H3K4me3 in heart but not liver by a Quantitative Trait Locus (QTL) on rat Chr 3 (Rintisch et al. 2014). Additionally, human studies are starting to identify *trans*-acting factors influencing DNA methylation and gene expression. One study using lymphocytes from 1,748 individuals (Lemire et al. 2015) found 1,657 trans-regulated sites scattered across the genome, while another study (Shi et al. 2014) examined the DNA methylome from 210 normal lung tissue samples, finding 373 QTL controlling 585 of 33,456 CpG sites. Finally, a recent comprehensive survey of 44 human tissue found *trans*-regulation of gene expression in 18 tissue, with testis having the most *trans*-acting expression QTL (Consortium 2017).

Here, we have employed the power and precision of mouse genetics to explore natural genetic variation in *trans*-regulation of the mammalian epigenetic landscape in two inbred laboratory strains of *Mus musculus domesticus*: C57BL/6J (B6) and DBA/2J (D2). B6 and D2 were the founder strains for a genetic panel of well over 100 Recombinant Inbred (RI) lines, which were created by intercrossing B6 and D2 mice followed by inbreeding the progeny to create the BXD genetic reference panel in which each line is a homozygous mosaic of the founder genomes (Taylor et al. 1973; Peirce et al. 2004). Panels of BXD lines have been used to facilitate mapping the genetic determinants of multiple phenotypes (Li et al. 2018).

As a model of epigenetic regulation, we focused on H3K4me3, a sensitive indicator of active chromatin (Santos-Rosa et al. 2002) in male germ cells. These cells provided the unique advantage that H3K4me3 sites are deposited by at least two independent enzyme systems, allowing us to distinguish between genetic variation specific to one of the enzyme systems and variation in underlying regulatory mechanisms that affect both. One system is the set of ubiquitous histone methyltransferases and demethylases that function in all cells to metabolize H3K4me3 at promoters and enhancers. The other is the meiosis-specific system placing H3K4me3 at genetic recombination hotspots. There, H3K4me3 is deposited by the histone methyltransferase PRDM9 (Baudat et al. 2010; Myers et al. 2010; Parvanov et al. 2010) and displaced when meiotic DNA double strand breaks are subsequently repaired (Sun et al. 2015). Importantly for this study, B6 and D2 mice share the same *Prdm9* allele, allowing the use of the BXD RI lines to identify genetic modifiers of hotspot H3K4me3 levels that act independently of PRDM9.

Carrying out ChIP-Seq for H3K4me3 in male germ cells we found quantitative differences between B6 and D2 mice. We identified six major *trans*-acting histone QTL (hQTL), located on Chrs 4, 7, 12 and 13, controlling the majority of *trans*-regulated H3K4me3 peaks. Additionally, by measuring H3K4me3 in embryonic stem cells (ESCs) and terminally differentiated hepatocytes and cardiomyocytes we find that *trans*-regulation of the epigenome is cell-type specific. In germ cells hQTL do not coincide with any of the known writers or erasers of histone H3K4 methylation and they affect both recombination hotspots and other functional elements. In F1 hybrids, low levels of methylation is the dominant phenotype, suggesting that hQTL genes suppress methylation. The locational specificity, ability to reduce active histone modifications, and lack of overlap with known enzymes of H3K4me3 metabolism, suggest that hQTL encode proteins that recruit chromatin repressing complexes. In accord with this hypothesis, hQTL regions contain KRAB- and BTB-zinc-finger proteins (ZFPs).

Taken together, these results indicate that a significant fraction of the epigenome is subject to *trans*-regulation by genes whose products bind at specific genomic locations to regulate the chromatin landscape.

## RESULTS

### Genetic variation influences histone modification levels

To obtain a genome-wide view of the similarities and differences between B6 and D2 mice in H3K4me3 sites, we performed nucleosome resolution ChIP-seq for H3K4me3 from germ cells of juvenile B6 and D2 inbred mice. Three biological replicates were collected from each strain by pooling germ cells from 3-5 mice for each replicate (Supplemental Table S1). Intra-strain variation between replicates was consistently low (Pearson correlation coefficient *r* = 0.92-0.98), while variation between strains was consistently greater (*r* = 0.80-0.88), suggesting genetic control of the chromatin state (Fig. 1A).

**Figure 1.**
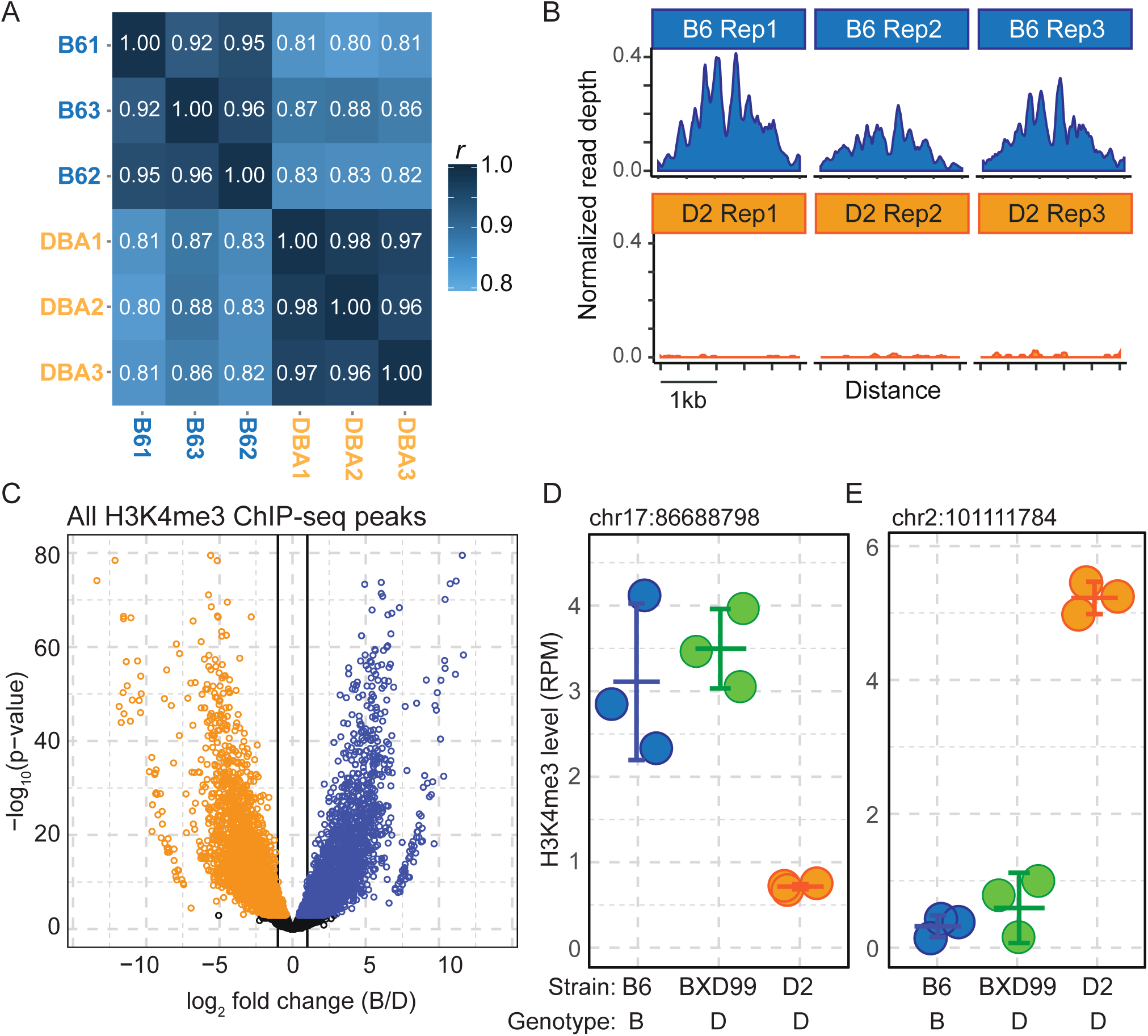
Trans-regulation of H3K4me3 levels in male germ cells. *(A)* Heat map of the Pearson correlation coefficients of H3K4me3 level between biological replicates of male germ cells isolated from B6 and D2 mice. *(B)* Coverage profile for a representative locus showing reproducibility of normalized (RPM) ChIP-seq read depth for biological replicates of a single H3K4me3 peak. *(C)* Volcano plot of differential H3K4me3 levels between B6 (blue) and D2 (orange) for all peaks (colored points represent peaks with significant difference after p-value adjustment, FDR < 0.05). *(D)* Example of an H3K4me3 peak under *trans*-regulation. This peak has high H3K4me3 level in B6 and low in D2; however, the BXD99 strain, which is genotypically D2 at this locus, has high levels of H3K4me3 similar to B6. Dots represent biological replicates for indicated strains. Error bars show mean and standard deviation. The local genotype at the H3K4me3 peak is shown below the plot, the genomic position of the H3K4me3 peak is above. *(E)* Similar to D showing an H3K4me3 peak with high H3K4me3 levels in the D2 background but low levels in BXD99, which is genotypically D2 at the locus.

In all, we identified 67,110 H3K4me3 peaks between the two strains. Among these peaks, 8,319 (12%) showed significant differences between strains (FDR adjusted p-value < 0.01), with approximately half having greater methylation in either B6 or D2. The level of differential activity varied considerably, from subtle changes in H3K4me3 level to sites that were virtually strain specific (Fig. 1B and C).

### Testing for *trans* control of epigenetic changes

We tested for *trans*-regulation of chromatin features, by exploiting the mosaic genetic structure of BXD lines. The location of each H3K4me3 peak identified in an RI line lies within a large, homozygous region, allowing us to identify the genetic origin of each site unambiguously. We defined an H3K4me3 peak as subject to *trans* control if its phenotype in an RI line differed significantly (p-adj <0.01) from the phenotype of the parental strain with the same genotype at that site (Fig. 1D and E). While this test is statistically quite robust, it underestimates the number of *trans*-regulated sites by at least 50% because an H3K4me3 peak is only detected as subject to *trans* control when its putative *trans*-acting hQTL has the opposite genotype from the H3K4me3 site. Applying this test to 3 biological replicates of RI line BXD99, we found that among the 8,319 peaks that were significantly different in the parents, 1,172 were *trans*-regulated in BXD99, indicating that at least a fourth of the differentially modified H3K4me3 sites are under *trans* control. For example, the H3K4me3 level of the peak at Chr 17 86.7 Mb (Fig. 1D) is low in the D2 parent and high in B6. The level of H3K4me3 at this site is also high in BXD99, even though the location of the peak in BXD99 places it in a region that is unequivocally homozygous D2. Similarly, we identified peaks with high H3K4me3 in the D2 parent, but low in BXD99, even though they were genetically D2 at the H3K4me3 locus (Fig. 1E).

### Linkage mapping in RI lines identifies multiple *trans*-acting QTL regulating chromatin

We mapped the genomic locations of the loci controlling H3K4me3 levels by performing H3K4me3 ChIP-seq in male germ cells of 60 BXD RI lines. To facilitate genetic mapping, we derived *de novo* genotypes using the ChIP sequencing data (Supplemental Fig. S1 and S2). To reduce mapping bias and improve quantification, we developed a custom alignment strategy accounting for known variation between B6 and D2 (see methods for details).

Using read depth for each peak as a quantitative phenotype, we genetically mapped the location of the hQTL controlling each H3K4me3 peak. For example, the level of H3K4me3 at the Chr2 10.1 Mb site, previously identified as under *trans* control in BXD99 (Fig. 1E), segregated as two distinct phenotypes in the BXD lines (Fig. 2A). Mapping the locus determining this phenotype identified a single hQTL on Chr 13 at 64.8 Mb (Fig. 2B). Having the D2 genotype at the Chr 13 hQTL resulted in a high level of H3K4me3 at the Chr 2 site (Fig. 2C), regardless of the local genotype on Chr 2, confirming the BXD test for *trans* control and identifying the location of this *trans*-regulator.

**Figure 2.**
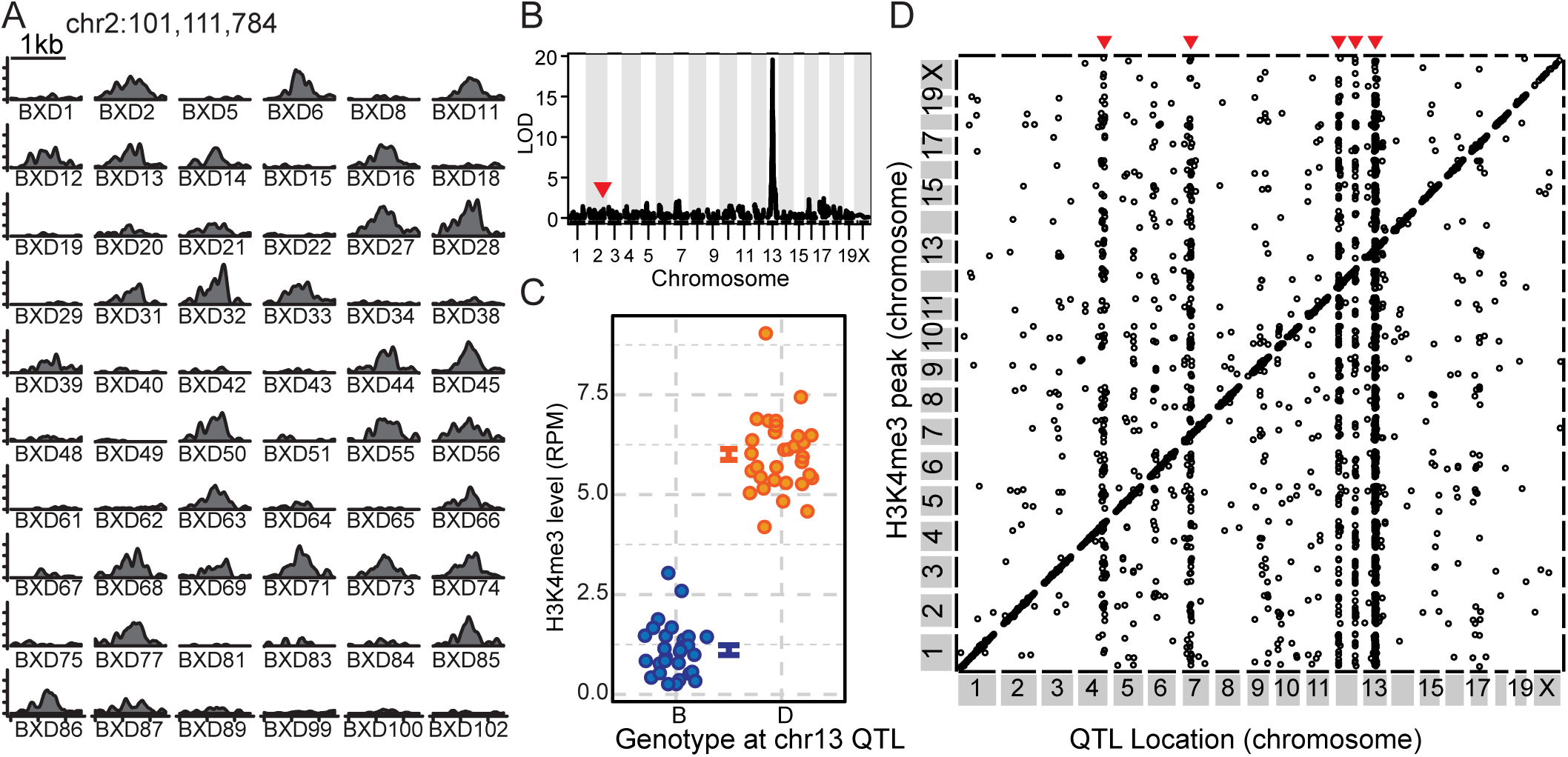
H3K4me3 levels are extensively regulated by local and distal hQTL. *(A)* Coverage profile for 60 BXD lines of H3K4me3 peak from Fig. 1E. *(B)* LOD score plotted against genetic map for H3K4me3 level shown in A. Location of the peak under regulation is indicated with red arrow. *(C)* H3K4me3 level plotted by genotype at the *trans*-hQTL marker on Chr 13 (error bars are standard error). *(D)* Points represent maximum LOD scores from single-hQTL scans with FDR < 0.05. The location of the H3K4me3 peak is plotted on the *y* axis and the location of the mapped hQTL on the *x* axis. Genetic effects acting locally (N=8,375) lie on the diagonal line. Distal acting *trans-*hQTL (N=1,867) lie off the diagonal; *trans*-hQTL impacting multiple loci appear as vertical bands (red arrows).

Overall, we identified 10,242 H3K4me3 peaks under genetic control in the BXD RI lines at a genome wide FDR < 0.05 after permutation testing (Fig. 2D, Supplemental Table S3). Among these, 5,160 had been detected earlier as statistically different between the B6 and D2 parents. A likely explanation for the increased sensitivity of RI lines is that testing 60 RI lines, rather than only 3 parental replicates, greatly increased our statistical ability to detect small phenotypic differences.

Among the 10,242 H3K4me3 peaks under genetic control, 1,867 were regulated in *trans* (FDR < 0.5) by hQTL that were on a different chromosome. This conservative definition of *trans* activity precluded any possibility of misidentifying *cis* activity at a long distance. Importantly, each *trans*-hQTL is highly specific, regulating a cohort of H3K4me3 sites. Among these *trans*-regulated H3K4me3 peaks, 1,335 (72%) are collectively controlled by 6 major hQTL on chromosomes 4, 7, 12, and 13 (Fig. 2D). The strongest hQTL, located on Chr 13, accounts for 41% of all *trans*-regulated sites.

### Epigenetic network analyses

We identified the same set of major hQTL by applying an iterative weighted correlation network analysis (iterativeWGCNA) (Langfelder and Horvath 2008; Greenfest-Allen et al. 2017) to maximize separation between co-regulated features, a method previously used for identifying refined gene clusters (modules). We identified 1,224 H3K4me3 peaks that grouped into 14 modules by their correlated levels of H3K4me3 modification (Supplemental Table S4). We mapped the loci controlling each module by combining the data for the peaks in each module into a single eigengene. All modules mapped to loci that overlapped the location of one of the *trans*-acting hQTL apparent in Fig. 2D, including Chrs 4, 7, 12, and 13 (Supplemental Table S4). Of the 14 modules, 9 map to Chr 13, representing 73% (n = 897) of all module peaks, with four modules having high phenotypes when D2 at the hQTL and five having high phenotypes when B6, suggesting compound genetic regulation at this locus. Furthermore, network analysis suggested that chromosome 7 has two distinct, but closely neighboring hQTL. These data provide independent confirmation of the six major hQTL, and show that individual hQTL regulate distinct cohorts of H3K4me3 peaks.

Although four of the major hQTL modules (mapping to Chrs 4, 7b, 12a and 13) act independently, with each module controlling a unique cohort of H3K4me3 sites, the network analysis showed that module 3, which maps to both Chr 7a and 12b, acts jointly on a distinct set of 162 H3K4me3 sites (Supplemental Table S4). Epistasis analysis did not detect interactions between 7a and 12b, suggesting that their effects are additive (data not shown). We conclude that hQTL 7a and 12b cooperate to regulate a single cohort of H3K4me3 sites.

### *Trans*-regulation of H3K4me3 is cell-type specific

Germ cells in the above study represent a mixture of cell-types representing both spermatagonial stem cells and differentiating meiotic cells. To reduce cellular complexity we applied the RI line test for *trans*-regulation of H3K4me3 to three distinct populations of highly enriched cell types: ES cells, hepatocytes, and cardiomyocytes (Fig. 3A). For each cell type, we performed H3K4me3 ChIP in a minimum of three biological replicates from B6 and D2 mice as well as from two RI lines, BXD75 and BXD87 (Supplemental Fig. S3). These data revealed hundreds of peaks under *trans*-regulation in each cell type. Using two different BXD lines provided internal controls for both reproducibility and specificity. For example, the H3K4me3 peak located on chr1 17.7 Mb in ES cells, which has high methylation in B6 and low methylation in D2 was identified in both BXD75 and BXD87 as under *trans*-regulation (Fig. 3B). Additionally, we found the expected ~50% overlap between the two BXD lines when we considered all predicted *trans*-regulated peaks (Supplemental Table S5). Very importantly, while all three cell types shared a considerable percentage of their total H3K4me3 sites, virtually all *trans*-regulated peaks were tissue-specific (Fig. 3C). Even though many H3K4me3 sites are found in multiple tissues (Fig. 3C brackets), *trans*-regulation of those peaks only occurred in a single cell type. Combining the results from germ cells, ES cells, hepatocytes and cardiomyocytes, we conclude that an extensive chromatin regulatory system operates in a cell-type specific manner to modulate H3K4me3 in *trans*.

**Figure 3.**
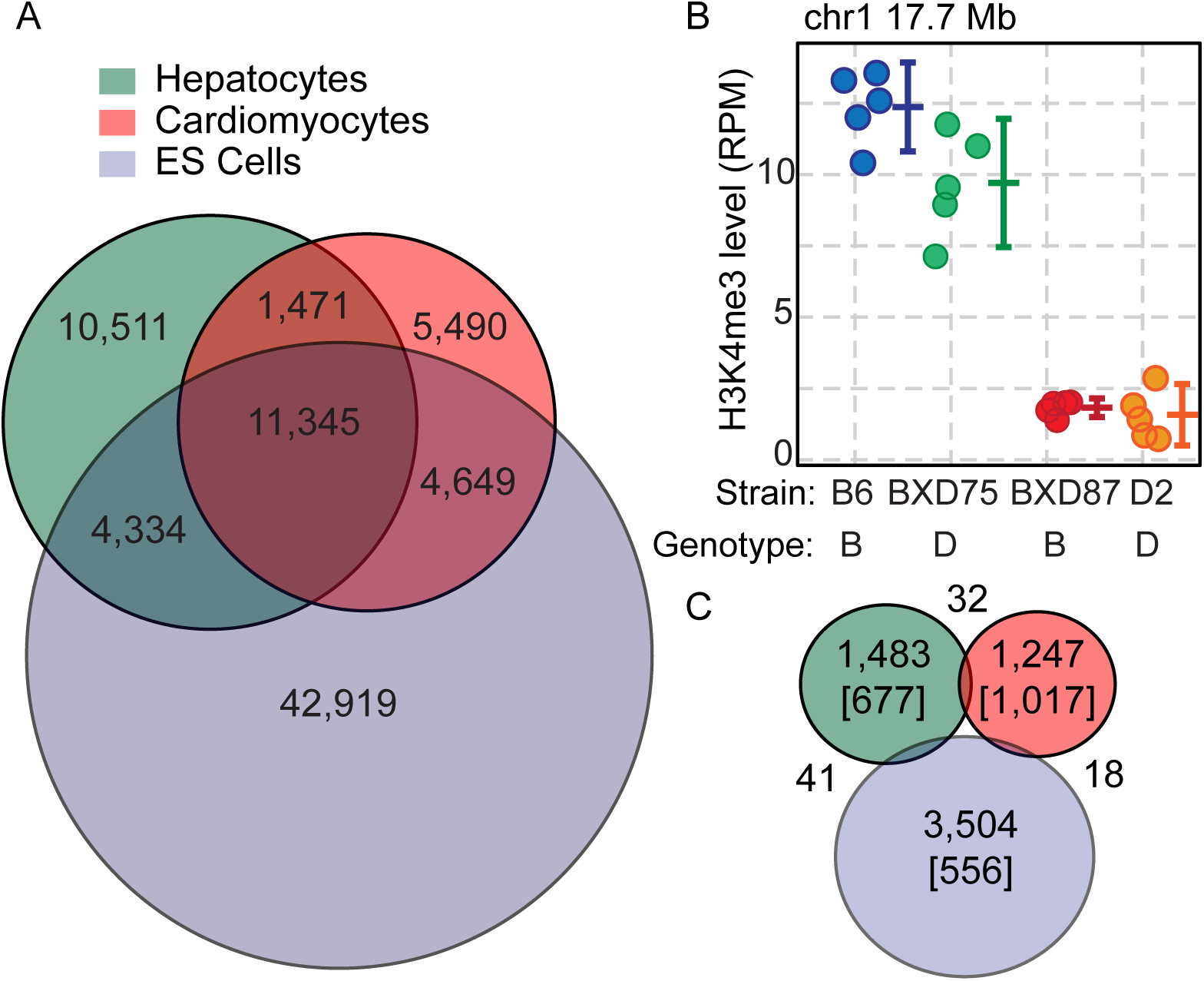
Trans-regulation of H3K4me3 is cell-type specific. *(A)* Venn diagram showing the number and relationship of all observed H3K4me3 sites between hepatocytes, cardiomyocytes, and embryonic stem cells from B6 and D2 parents. *(B)* Example H3K4me3 site predicted to be under *trans*-regulation in embryonic stem cells. Both the strain and genotype at the H3K4me3 site is indicated under the graph (error bars represent standard deviation). *(C)* Venn diagram of the number and relationship of predicted *trans*-regulated H3K4me3 sites across three cell types. Numbers in brackets indicate peaks that, while being uniquely subject to *trans*-regulation within the indicated cell type, are present in at least one of the other cell types.

### Germ cell hQTL dominantly suppress H3K4me3

To determine whether hQTL alleles act additively or dominantly in germ cells, we compared H3K4me3 levels from germ cells in heterozygous (B6xD2)F1 hybrids with those in the two parental strains. A *cis*-acting local variant is expected to control each chromosome independently in an F1 heterozygote, resulting in additive inheritance, with the phenotype of the F1 intermediate between B6 and D2 (Fig. 4A). In contrast, a *trans*-acting hQTL that either induces or suppresses H3K4me3 levels would likely show a dominant phenotype in the F1 that would be closer to one of the parents. For example, the D2 allele of the Chr 13 hQTL dominantly suppresses H3K4 trimethylation at site on Chr 5 in *trans* (Fig. 4B and C).

**Figure 4.**
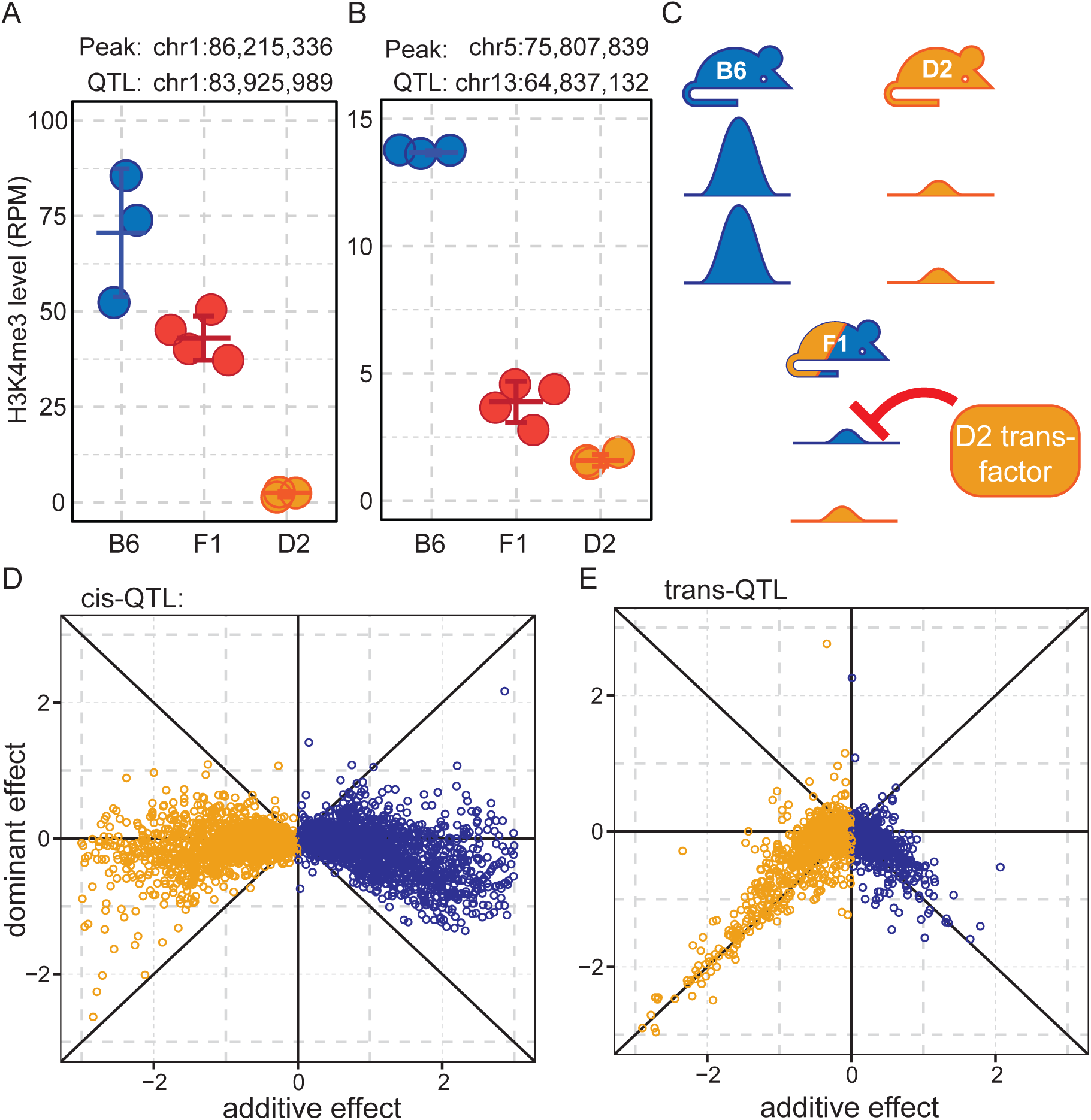
Trans-hQTL suppress H3K4me3 levels in F1 hybrids. *(A)* Example of a local-hQTL with additive inheritance in F1 hybrids. *(B)* Example of a distal-hQTL showing dominant repression of H3K4me3 activity in F1 hybrids, similar to the D2 parent. *(C)* Model illustrating the dominant effect in F1 hybrids. A D2-derived *trans*-acting factor results in reduced H3K4me3 on both chromosomes in the heterozygous state. *(D)* All *cis*-hQTL with LOD > 20 showing most loci exhibit additive inheritance. *(E)* All *trans*-hQTL showing the general dominant repression of H3K4me3 levels.

To generalize these observations, we adopted the method of Tian et. al. (Tian et al. 2016) in which the measure of dominance (d), defined as the F1 minus the parental average, is plotted against the size of the additive effect (a), which is defined as half the distance between the mean of the two parents. In this formulation d is zero when there is no dominance, i.e. additive inheritance; d is positive when high methylation is dominant, and negative when low methylation is dominant. As expected, additive inheritance is generally seen at *cis*-QTL (LOD score > 20, Fig. 4D). When plotting all *trans*-acting hQTL together, regardless of whether the H3K4me3 site is high in B6 or D2, we observe the trend expected for fully dominant reduction in methylation (Fig. 4E). We conclude that the *trans*-hQTL we observe in germ cells all act by suppressing H3K4me3.

### hQTL loci can be genetically compound

Many of the major hQTL are defined by H3K4me3 peaks whose QTL map to adjacent genetic intervals along the chromosome. This, and the fact that multiple network modules can be assigned to the same regions, suggest the possibility that some hQTL are compound, containing more than one causal gene. To refine the location of candidate regions we took advantage of the homozygous genetic structure of the BXD genomes, focusing on the RI lines with crossovers in the QTL regions and the H3K4me3 peaks that showed strong difference in phenotype between B6 and D2 (for example Fig 2A-C). When individually mapped, 613 peaks (FDR < 0.05) regulated by Chr 13 are associated with 6 consecutive intervals across an ~8 Mb region (Supplemental Table S3). The RI lines that are most informative about a causal gene’s exact location are the lines with crossovers that divide the QTL region in question. Out of the 60 strains used in this study 7 have recombination breakpoints across this region (Supplemental Figure S4). Plotting the H3K4me3 level for several informative peaks based on the genotype at each of the 6 intervals shows a clear split in *trans*-regulation across this region, defined by a crossover in BXD71, and suggests multiple closely spaced hQTL. In the case of Chr 13, this analysis found hQTL in at least two genetically distinct regions in close proximity: Mb 60,418,152-65,890,145 and 65,890,145-69,026,189 (Supplemental Fig. S4). Repeating this analysis on other hQTL regions, resulted in a suggestive split at the Chr 4 hQTL (Mb 143,302,894-144,911,401 and 144,911,401-148,865,285) determined by a crossover in BXD42.

### *Trans*-regulation of chromatin involves an array of genomic regulatory features

The chromatin regulatory genes within the hQTL differ from traditional transcription factors in that they are capable of impacting all classes of H3K4me3 sites in male germ cells – promoters, enhancers, insulators, transposons, and recombination hotspots, as well as sites of unknown function (Supplemental Fig. S5). Overall, *trans*-QTL regulate all classes of functional elements, although individual modules differ in the proportions of functional elements they influence (Supplemental Fig. S5). In particular, the hQTL on Chr 7b and Chr 12a (represented by modules 2 and 5) are enriched for hotspot sequences, while hQTL 7a and 12b (module 3), which act on an overlapping set of H3K4me3 peaks, are virtually devoid of hotspots. Of the 9 modules that map to Chr 13, most are depleted for hotspots, with the exception module 8. While many individual modules are depleted for promoters and genic regions, module 1 on Chr 4 is enriched. Our male germ cell preparations include both spermatogonial stem cells and spermatocytes; given the evidence of cell-type specificity cited above, these differences among hQTL enrichments suggest that they may act at different stages of meiosis.

### hQTL influence cellular function

A unique property of meiotic cells offered a direct test of whether the hQTL have functional consequences. The meiotic DSBs that give rise to genetic recombination occur following activation of hotspots by PRDM9, in part through local deposition of H3K4me3 and H3K36me3 (Smagulova et al. 2011; Baker et al. 2014; Lange et al. 2016). Accordingly, we tested whether hQTL regulation of H3K4me3 levels at recombination hotspots influences the subsequent likelihood of DSB formation by comparing H3K4me3 levels in BXD mice to DMC1 levels in the founders (Fig. 5A and B).

**Figure 5.**
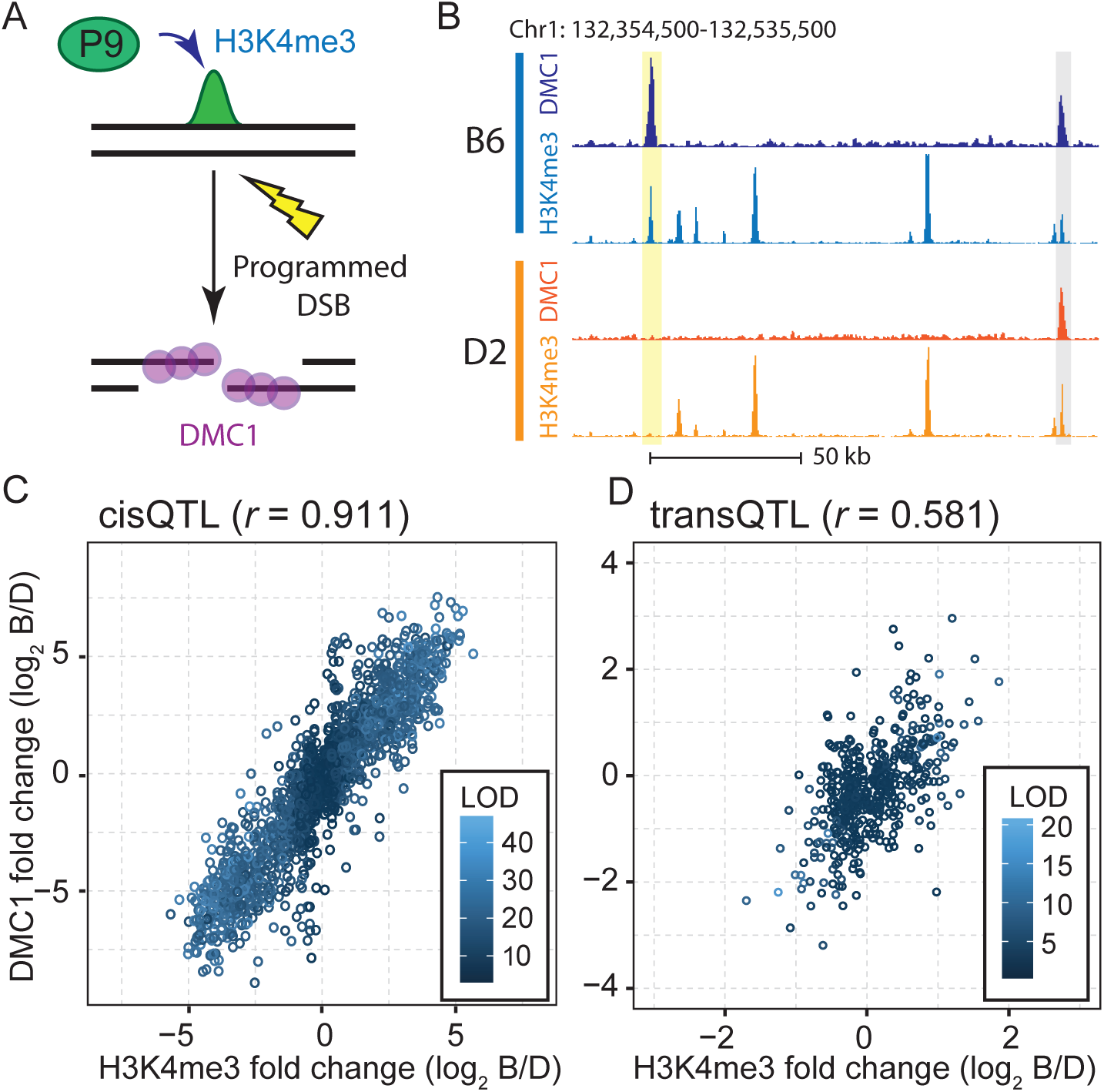
hQTL control DSB frequency at recombination hotspots. *(A)* Illustration of events leading up to meiotic DSBs. PRDM9 (P9) binds DNA and locally trimethylates H3K4, leading to targeting of meiotic DSB machinery. The DSB is resected and single-stranded DNA ends are coated with the meiosis-specific recombinase DMC1. *(B)* Coverage profile showing H3K4me3 and DMC1 ssDNA levels; yellow – example recombination hotspot that is active in B6 but not D2; grey – example hotspot found in both strains. *(C)* Scatterplot of the log2 ratio of B6/D2 H3K4me3 levels found in BXD strains versus parental B6/D2 DMC1 levels at hotspots for all *cis*-hQTL, showing Pearson correlation (*r*). *(D)* Similar to C showing results for all hotspots under regulation by *trans*-hQTL.

Doing so, we found a clear correlation between hQTL determined changes in H3K4me3 levels and the consequent changes in DSB frequency at both *cis*- and *trans*-hQTL. This was true for both *cis*- and *trans*-acting hQTL (Fig. 5C and D). For *cis*-acting hQTL, the differences in H3K4me3 levels account for nearly 83% of the variance in DSB frequency; it was less so for *trans*-acting QTL, which are likely subject to additional genetic variation. We conclude that hQTL modulation of PRDM9-dependent H3K4me3 influences the subsequent likelihood of DSB formation at hotspots. In effect, chromatin regulatory loci influence the recombination landscape and thus inheritance patterns between generations.

Additional evidence that hQTL are functionally distinct is supported by the observation that the H3K4me3 peaks found in several modules contain distinct sets of transcription factor (TF) binding sites. For example, module 3 (Chrs 7a and 12b) is highly enriched for the YY1 motif (p-value < 8.98×10^−29^) and *Yy1* is located on Chr 12, two Mb from the Chr 12b critical region. YY1 is an architectural protein that bridges the physical interactions between cell-type specific enhancers and promoters (Beagan et al. 2017; Weintraub et al. 2017), suggesting that the H3K4me3 peaks regulated by module 3 could be meiosis-specific functional elements. Additionally, several modules are enriched for TF motifs that are predicted to form regulatory networks in meiotic cells (Ball et al. 2016). For example, module 9 is enriched for the well-known TF retinoic acid receptor *Rarg* (p-value < 2.19×10^−5^), which is upregulated in preleptotene spermatocytes, a meiotic substage abundantly present in our male germ cell preparations. Module 1 is enriched for the *Zbtb33* motif (p-value < 5.73×10^−3^), a TF expressed later in meiosis, and is also enriched for H3K4me3 peaks that overlap promoters and known testis enhancers (Supplemental Table S4). Finally, modules 3 and 11 are enriched for *Zfp42* motifs to different extents (p-values < 1.46×10^−20^ and 6.93×10^−3^ respectively). *Zfp42* is expressed exclusively in pluripotent embryonic stem cells and male germ cells (Rogers et al. 1991); it is a known marker for pluripotency and functions to repress transposable elements (Schoorlemmer et al. 2014). Together, these data show that different modules are enriched for different transcription factor binding motifs, suggesting these modules may represent functionally related sets of H3K4me3 peaks that interact with known meiotic TFs.

### *Cis* regulation of H3K4me3 level

In addition to *trans*-acting hQTL, 8,375 hQTL map close to the location of the H3K4me3 peak they control. At each of these peaks, the difference in H3K4me3 level between BXD lines whose genotype at the peak is B6 versus those with a D2 genotype are highly correlated with the differences in H3K4me3 level between the B6 and D2 parents (Fig. 6A). Defining *cis*-regulated H3K4me3 sites as those whose hQTL was within 10 Mb, we found an increased frequency of sequence variants concentrated near the center of H3K4me3 peaks compared to *trans*-regulated sites (Fig. 6B). *Cis*-hQTL were also significantly enriched for recombination hotspots compared to all peaks (Supplemental Fig. S5); hotspots are known to undergo selection for motif-disrupting mutations in their PRDM9 binding sites (Myers et al. 2010; Baker et al. 2015). An example is the locally controlled PRDM9-dependent H3K4me3 peak shown in Fig. 1B. The nucleosome depleted region where PRDM9 binds in this H3K4me3 peak (Baker et al. 2014) contains a match to the PRDM9 consensus sequence in the B6 genome, but there are two potentially disrupting SNPs in the motif in the D2 genome (Fig. 6C and D). Extending this observation, we found an increase in single nucleotide variants at PRDM9 binding motifs (Fig. 6E). Out of the 1,516 recombination hotspots with a *cis*-hQTL, 61% (n = 929) have a variant within the PRDM9 binding site (Fig. 6F, 36 base pair motif ± 4 bp) (Walker et al. 2015). That 39% of *cis*-regulated hotspots lack variants within their PRDM9 binding motif indicates the existence of additional mechanisms for *cis* control of hotspots. An appealing possibility is that these hotspots are in the vicinity of genetically variable DNA binding sites for other *trans*-regulatory proteins that do not differ genetically between B6 and D2.

**Figure 6.**
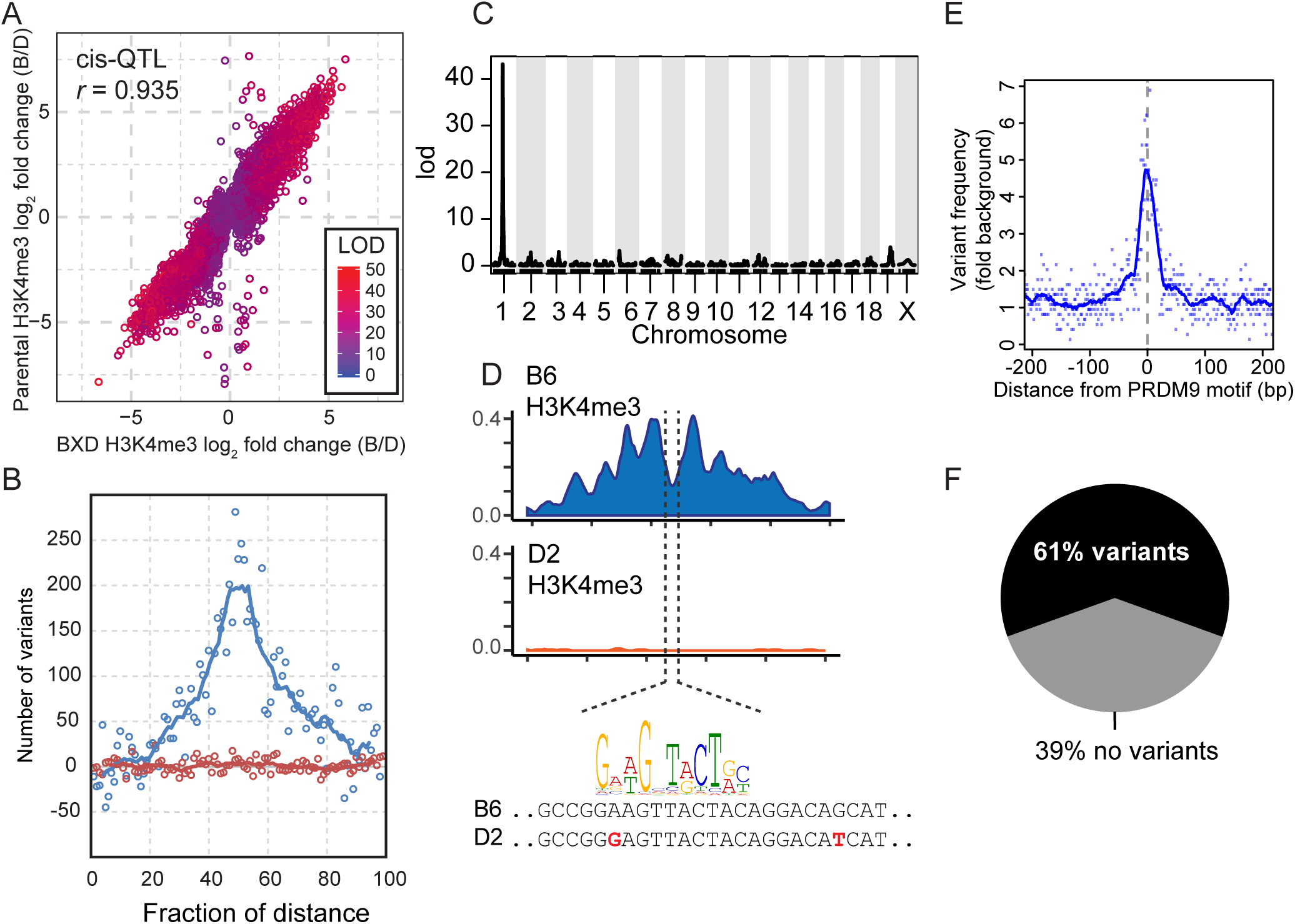
Local cis-hQTL are enriched for genetic variants within the H3K4me3 peak. *(A)* Correlation of the log fold change (B6/D2) between the founders and BXD lines for H3K4me3 levels at local-hQTL (points shaded by LOD score). *(B)* Number of background-subtracted genomic variants between B6 and D2 (SNPs and short indels) as a fraction of the distance across H3K4me3 peaks (blue – local-QTL, red – distal-QTL, solid lines – 5 base pair moving average). *(C)* LOD plot for local-QTL for an example recombination hotspot on distal Chr 1. *(D) upper* – Coverage profile of H3K4me3 level (loess-smoothed RPM) for hotspot shown in C for both genetic backgrounds. *lower* – DNA sequence from nucleosome depleted center of recombination hotspot showing variants between strains in red. *Prdm9^Dom2^* motif is indicated above best matching DNA sequence. *(E)* Enrichment of genomic variants between B6 and D2 plotted against distance from *Prdm9* motif for all local-QTL at recombination hotspots (solid line – 10 base pair moving average). *(F)* Percent of recombination hotspot local-QTL that have known genetic variants within the *Prdm9* binding site.

## DISSCUSSION

The totality of our data imply the existence of an extensive set of regulatory genes controlling chromatin modifications that are likely to be distinct from the enzymes that establish or remove methylation at H3K4me3. These genes, represented by *trans*-acting hQTL, control specific cohorts of chromatin sites, and act dominantly to suppress trimethylation of H3K4. Testing four diverse cell populations we find cell-type specific *trans*-regulation of H3K4me3. Several of the hQTL are genetically compound, suggesting they contain multiple regulatory genes. Two independent lines of evidence suggest that these chromatin regulatory genes have functional significance. They affect the likelihood that recombination hotspots can acquire the DSBs that give rise to genetic crossing over, and cohorts of H3K4me3 sites controlled by individual hQTL contain distinct sets of transcription factor binding sites.

The broad extent of chromatin regulation makes it likely that the activities of many chromatin regulatory genes are themselves influenced by other chromatin regulatory genes, creating a series of regulatory loops that interact to determine the overall chromatin state of a cell. In this case, differentiation can be viewed as a progression of epigenetic states determined by feedback driven, consequential changes in expression of chromatin regulatory genes, until a final, stable homeostatic state is reached. In essence, this is simply putting a molecular face to Waddington’s original seminal concept of epigenetic canalization (Waddington 1957).

### Distinctions among hQTL

Male germ cell hQTL modules differ among themselves in several respects. Module 3 is unique in that both loci act on the same cohort of H3K4me3 peaks, it does not modulate recombination hotspots at all, and is unidirectional in the sense that nearly all H3K4me3 peaks under regulation are high when the same allele of the hQTL is present. In contrast bidirectional hQTL are mixtures of high and low H3K4me3 peaks when the B6 (or D2) allele is present. In contrast to module 3, the other major hQTL modules are bidirectional, affect hotspots, and each controls a unique cohorts of H3K4me3 peaks. Bidirectionality can be explained if the alleles of a chromatin regulatory gene differ in their DNA binding specificity, act at different stages of meiosis, or if the QTL is genetically compound. Cohort sharing and lack of influence on hotspots by module 3 can be explained if the gene products function together in a pathway that is only expressed at a stage of meiosis when recombination hotspots are not present, for example in spermatogonia.

### Potential candidate genes

It is unlikely that the genes underlying the hQTL are enzymes that write or erase H3K4me3. First, many of the hQTL, including hQTL 4, 7b, 12a and 13, regulate both recombination hotspots and conventional promoter and enhancer sites, locations where H3K4me3 is metabolized by independent enzyme systems. Second, all of the hQTL show great locational specificity, but the known writers or erasers of H3K4me3 are not known to possess DNA binding activity on their own. And third, the locations of our major hQTL did not coincide with the locations of any of the genes encoding these enzymes. H3K4 trimethylation is catalyzed by six SET domain proteins: SET1A, SET1B, and MLL1-4, which nucleate the COMPASS complexes containing ASH2L, RBBP5, WDR5, and DPY30 plus 2-4 additional proteins chosen from among a battery of possible accessory proteins (CFP1, WDR82, BOD1/1L, HCF1/2, NCOA6, MENIN, KDM6A, PSIP1, PAXIP1, and PA1) (Gu and Lee 2013). H3K4me3 at transcriptional regulatory sites can also be formed by the action of SMYD1, 2 and 3, ASH1L, SET7/9, and SETMAR. In addition to these writers, there are several known H3K4 demethylases of the LSD and JARID families (Gu and Lee 2013). Importantly, none of the genes encoding these proteins coincide with the locations of the six major QTL regions.

While we cannot rule out that the hQTL may modulate the abundance or activity of one of the above genes, this type of regulation does not provide an explanation for the observed locational specificities of our hQTL. Instead the properties of the identified hQTL, ability to suppress H3K4 methylation, and the physical location of the hQTL all make zinc finger proteins (ZFPs), particularly KRAB-ZFPs, attractive candidates. In germ cells, there is a considerable coincidence in location between the observed hQTL and ZFPs. Most of the hundreds of ZFP genes occur in a small number of tightly linked clusters (Walker et al. 2015; Kauzlaric et al. 2017), and the major hQTL on Chrs 4, 7b, 12a, and 13 all overlap such clusters, while the hQTL at Chr 7a and 12b each overlap a single zinc finger gene. Moreover, in the case of the Chr 13 QTL, which is compound, each of the sub-regions overlaps a different ZFP cluster. The presence of multiple ZFPs within an hQTL could help to explain bidirectionality as different ZFPs would impact different sites in the B6 and D2 genomes. Finally, the Regulator of sex limitation genes (*Rsl1* and *Rsl2*) provide precedents. These KRAB-ZFPs repress expression of a variety of genes in female liver, but have no effect in kidney (Krebs et al. 2003). At one target gene, *Slp* (Sex limited protein), RSL1 binds at an LTR-based enhancer upstream of the *Slp* promoter, where suppression correlates with methylation of CpG dinucleotides (Krebs et al. 2012). Incidentally, *Rsl1* and *Rsl2* are located in the distal Chr 13 hQTL ZFP cluster, although we do not yet have sufficient evidence to rule these genes in or out as candidates in germ cells.

Zinc finger proteins make up the single largest class of genes in mammalian genomes (Ecco et al. 2017), and many possess secondary domains that nucleate chromatin modifying complexes capable of closing chromatin. This is particularly true of KRAB-ZFPs, of which there are several hundred in both mice and humans (Ecco et al. 2017), and BTB-ZFPs with around 50 identified in both species (Stogios et al. 2005). Their zinc fingers provide DNA binding specificity, accounting for the locational specificity of H3K4me3 peaks regulated by hQTL, and their KRAB and BTB domains generally recruit chromatin repressing complexes to these specific genomic locations, accounting for their dominant suppression of H3K4me3. KRAB domains frequently bind TRIM28/KAP1 (Friedman et al. 1996), leading to deacetylation of H3K27 and trimethylation of H3K9 (Ecco et al. 2017), two reactions that serve to repress or close chromatin. BTB domains can close chromatin by associating with histone deacetylase complexes including N-CoR and SMRT (Huynh and Bardwell 1998). Together these chromatin modifying features of ZFPs combine to establish cell-type specific gene regulatory networks (Krebs et al. 2012; Ecco et al. 2017; Yang et al. 2017). Furthermore, a recent report found a correlation between reduced levels of meiotic recombination and proximity to KRAB-ZFP binding sequences (Altemose et al. 2017). Combined, these properties of our identified hQTL suggest a model in which DNA-binding factors guide chromatin repressing complexes to specific genomic locations in a tissue-specific manner, thereby establishing major features of the epigenome.

### Testing for *trans*-regulation of genomic features

Classically, testing for *trans*-regulation relies on asking whether chromosomal sites that are regulated differently in two parents behave the same in the heterozygous background of an F1 hybrid (Wittkopp et al. 2008). In attempting to apply this test to B6 and D2 mice, we found it limited by the paucity of informative SNPs between the two haplotypes. Instead, using the homozygous genome of an RI line, we can reliably conclude that a chromosomal site is subject to *trans*-regulation in if it’s molecular phenotype does not correspond to the genetic origin of the DNA at that site. If an H3K4me3 peak in a BXD line is genetically B6 in origin but no longer has the phenotype seen in B6 mice, we conclude that it is under *trans* control by one or more distal hQTL; similarly, for sites of D2 origin. In practice this advantage more than compensated for the 50% efficiency resulting from the fact that the test is only positive when the distal QTL has the opposite genotype of the regulated H3K4me3 site.

Our hQTL mapping provided an opportunity to confirm the utility and accuracy of this test. hQTL mapping in 60 RI lines identified 4,429 *cis*-hQTL that were also significantly different in the parental lines. Testing a single RI line, BXD99, over 95% (4,226) of H3K4me3 sites showed similar phenotypes to their parents of origin and therefore were correctly predicted to be under local regulation. Conversely, we identified 731 *trans*-hQTL in RI lines that showed significant parental differences and 43% of these were predicted as *trans* in BXD99, a value similar to the estimated 50% efficiency. Additionally, in the other three cell types tested, we see nearly 50% overlap in the identity of the H3K4me3 peaks predicted to be under *trans*-regulation between different BXD lines.

In summary, we have observed *trans*-regulation of the epigenome in four functionally very diverse cell types. ES cells are pluripotent, hepatocytes are capable of tissue regeneration and are endodermal in origin, cardiomyocytes are not capable of regeneration and are mesodermal in origin, and our germ cell preparations include a mixture of meiotic cell stages progressing from spermatogonial stem cells through early meiosis. Extrapolating these results, it seems likely that *trans*-regulation of the epigenome occurs in many mammalian cells, with significant implications for human health. An estimated 93% of mapped genetic variation impacting human health lies outside the coding regions of genes in functional elements (Maurano et al. 2012; Albert and Kruglyak 2015). Additionally, chromatin regulatory loci, such as those identified as hQTL here, have the potential to subtly shape the activity of functional elements in *trans* and therefore influence the manifestation of complex traits.

### Data Access

Raw sequencing data and processed files are available at the NCBI Gene Expression Omnibus (GEO; http://www.ncbi/nlm/nih.gov/geo/) available under accession number GSEXXXXX. DMC1 ChIP-seq data for B6 is available under accession number GSE108259.

## Acknowledgments

We thank members of the Baker, Paigen, Petkov, and Carter labs for their discussion of the data and manuscript. This work was assisted by JAX scientific services that are supported through NIH Cancer Core grant CA34196. BXD ES cell lines were kindly provided by Anne Czechanski and Laura Reinholdt, funded by the Special Mouse Strain Resource OD011102-18. Funding for the work was provided by NIGMS F32 GM101736 and The Jackson Laboratory start-up funds supporting C.L.B., and P01 GM099640 to K.P and G.W.C.

## Author Contributions

C.L.B., P.P., N.P., H.T., C.S., and D.R. performed experiments. C.L.B., M.W., S.A., G.A., B.J., and K.B performed computational data analyses. C.L.B. and K.P. contributed to experimental design. C.L.B., G.W.C., and K.P supervised the study. C.L.B. and K.P wrote the manuscript. All authors discussed the results and interpretation, read and approved the final manuscript.

**Disclosure Declaration** The authors declare no competing financial interests.

## METHODS

### Mice

C57BL/6J (stock number 000664), DBA/2J (stock number 000671), B6D2 F1/J hybrid (stock number 100006), and all BXD recombinant inbred mice were obtained from The Jackson Laboratory, Bar Harbor, USA. All animal experiments were approved by the Animal Care and Use Committee of The Jackson Laboratory (Summary #04008 and #16043).

### Tissue isolation

Testicular germ cell enrichment was performed on 14 day post-partum male mice as previously reported (Baker et al. 2014).

Individual low-passage mESCs were derived using protocols outlined in (Czechanski et al. 2014). Briefly, 6-8-week-old females are mated to stud males and checked each morning for plugs. Pregnant females are euthanized on embryonic day 3.5 and the uterine horn is flushed to remove embryos. Embryos are visualized under a dissecting microscope and blastocysts are transferred to 2i (2i:CHIR99021 and PD0325901) (Ying et al. 2008) serum-free media for outgrowth of the ICM. Blastocysts are allowed to hatch and attach to the feeder layer, and the resulting outgrowth is monitored daily and fed for 8-11 days. The emergent ESCs are disaggregated and passaged onto new MEF feeders. Cultures during this time are closely monitored for unusually rapid growth (potentially indicating karyotypic instability), signs of deterioration including vacuolated cytoplasm, detachment of cells from colonies and debris, and possible signs of contamination. Successful ESC cultures were maintained on MEF feeders in serum containing 2i media supplemented with Leukemia inhibiting factor (2i/LIF) (Kiyonari et al. 2010) to maintain high levels of NANOG expression, which indicates ground-state pluripotency (Ying et al. 2008; Czechanski et al. 2014). Prior to preparing ESCs for chromatin isolation, mESCs were enzymatically disassociated using trypisin and MEFs were removed by serial plating on gelatin-coated plates to which MEFs adsorb preferentially; for this, ESCs and MEFs are incubated in 2i/LIF media on fresh plates for 15 minutes to allow the larger MEFs to quickly attach to the plates. ESCs are aspirated and the plating procedure repeated once to further remove MEFs. ESCs were collected by centrifugation, resuspended in PBS, and crosslinked using formaldehyde.

For hepatocyte isolation and purification, livers from eight-week-old female mice were perfused using a modified EGTA–collagenase perfusion protocol (Neufeld 1997). All perfusions and hepatocyte purifications were done at the same time of the day to avoid possible circadian effects on any studied parameter. EGTA buffer was used to flush the blood out of the liver and start to digest the desmosomes connecting the liver cells. After 35ml of the 1x EGTA solution was passed through the liver, it was replaced with 7-10ml of 1x Leffert’s buffer to flush out the EGTA, which otherwise chelates the calcium ions necessary for collagenase activity in the next step when the liver is digested by prefusion with 25-50 mL of Liberase solution (~4.3 Wünsch units). After perfusion, the liver was removed from the abdominal cavity and passed through Nitex 80 μm nylon mesh, using extra ice-cold Leffert’s buffer with 0.02% CaCl2 and a rubber policeman. Hepatocytes were purified from the remaining cells by two consecutive centrifugations for 4 min, 50 x g each, leaving the other, smaller cell types in suspension. After each spin, the solution was decanted as waste, and the enriched cell pellet of hepatocytes was resuspended in 30 ml ice-cold Leffert’s buffer with 0.02% CaCl2. After the second centrifugation, the cell pellet contained >98.6% hepatocytes.

For cardiomyocyte isolation eight-week-old female mice were euthanized and the chest opened to expose the heart. The descending aorta and inferior vena cavae were cut and an EDTA buffer was injected into the apex of the right ventricle to flush the heart. The ascending aorta was clamped and the heart transferred to a petri dish and fixed by perfusion of EDTA buffer containing 4% formaldehyde via the left ventricle. The formaldehyde was quenched by perfusing the heart with 125mM glycine, and digested by perfusion with collagenase buffer. The ventricles were rent into smaller pieces, and triturated to complete cellular dissociation into a single-cell suspension. Cells were then filtered through a 100-um strainer to remove tissue fragments and centrifuged at a very low speed to obtain a highly enriched fraction of fixed cardiomyocytes.

### H3K4me3 Chromatin Immunoprecipitation

ChIP was performed using H3K4me3 antibody (EMD/Millipore, #07-473) on chromatin sheared enzymatically as previously reported (Baker et al. 2014). Briefly, after isolation of single-cell suspensions, cells are cross-linked with formaldehyde followed by enzymatic chromatin shearing using micrococcal nuclease (MNase). Magnetic Protein-G-coated beads (Dynabead, Life technologies) are preloaded with H3K4me3 antibody and incubated with MNase-digested nucleosomes. Beads are washed and DNA is eluted and deproteinated, and crosslinks reversed using glycine buffer.

DNA was prepared for high-throughput sequencing using either the Bioo Scientific’s NEXTflex ChIP-seq Kit without size selection for all germ cell data or the KAPA hyper kit (Kapa Biosystems) for mESC, cardiomyocytes, and hepatocytes. Library quality and size distribution were visualized using a Bioanalyzer (Agilent). All samples are sequenced in-house using either the Illumina^®^ HiSeq 2500 or 4000 platform.

### Data analysis and H3K4me3 quantification

All sequenced B6, D2, F1 and BXD H3K4me3 ChiP libraries, as well as all control input DNA samples, were aligned utilizing bwa version 0.7.9a (Li and Durbin 2009). B6 parental samples were aligned to the Genome Reference Consortium Mouse Build 38 (mm10) and D2 parental samples were aligned to the *de novo* REL-1509-assembly, including all unplaced scaffolds, from the Mouse Genomes Project (Yalcin et al. 2012).

To ensure that H3K4me3 peaks were properly quantified across divergent genomes, we began by building a comprehensive “peakome” representing all potential H3K4me3 peaks found in the two parents. H3K4me3 peaks for B6 and D2 were called independently, utilizing alignment data from three replicate samples and one DNA input sample. Reads were filtered for a mapq alignment metric of 60 and an alignment sequence having no indels present across the entire length of the sequencing read, typically 100 bp. H3K4me3 peaks were called utilizing MACS version 1.4.2 (Zhang et al. 2008) and peaks having a false discovery rate of <1% found in two out of three replicates were accepted. Final genomic intervals for each H3K4me3 peak for each strain were derived by merging the peaks from the corresponding replicate samples using bedtools (Quinlan and Hall 2010). To link syntenic regions between B6 and D2 assemblies, which each have their own coordinate system, sequences from these genomic intervals were aligned to their alternative genome using reciprocal BLAST. In some cases a sequence interval comprising an H3K4me3 peak in one strain aligned to multiple adjacent intervals in the alternative genome. If the sequences of these peaks in the alternate strain all fell within the boundaries of the single peak, they were merged. The boundaries of these merged peaks included the incorporated sequences from both strains. Because there are also H3K4me3 peaks that are strain-specific, these peaks were accepted if, and only if, the mapped interval had a unique sequence that was found in the proper syntentic order within the alternative genome lacking that H3K4me3 peak. The final combined "peakome" between B6 and D2 mice was created by selecting only peaks appropriately linked across each strain, assuring that each H3K4me3 peak reciprocally aligned to only one peak in the alternative genome after merging, and that all peaks were in the same order along the chromosomes in both genomes (Supplemental Table S2). Using the H3K4me3 peaks locations derived from the parental strains, final read counts for B6, D2, F1 hybrids, and BXDs were obtained by counting reads within the coordinate boundaries of the peakome intervals.

To improve mapping accuracy and utilize known sequence variation between strains, all BXDs and F1 hybrid samples were aligned separately to both the mm10 reference and the *de novo* D2 assembly. To reduce error in quantification of H3K4me3 levels due to genomic regions containing repetitive sequences, we removed reads with multiple alignments and retained reads with alignment metric of 60 that lacked small indels, which can often indicate mis-alignment. Subsequently, for each genomic interval in the peakome, final reads counts were summed for those that mapped uniquely to one of the assemblies along with those that mapped equally well to both B6 and D2 genome assemblies.

### Genetic map construction

During the course of QTL mapping we found occasional discrepancies between the phenotype of a given H3K4me3 peak and its assigned local genotype when using publicly available genetic maps (Supplemental Fig. S1). These were particularly apparent for *cis*-QTL with large LOD scores where the H3K4me3 levels show distinct segregation between B6 and D2. To correct for these discrepancies and improve genetic mapping, genotype maps were derived *de novo* by calling genotypes of mm10 aligned reads from BXD H3K4me3 ChIP-Seq samples used in this study at known SNP positions between B6 and D2 reported in Mouse Genomes Project database REL-1505 (Keane et al. 2011; Yalcin et al. 2011). The location of genetic crossovers were assumed to lie within the transition intervals between B6 and D2 genotypes in each BXD line. Transition intervals comprise the DNA sequence between the base-pair position of the SNP with the first detectable genotype switch and the previous SNP. In some cases transitions could be detected within individual recombination hotspot H3K4me3 peaks (Supplemental Fig. S2). Overall, the sizes of the transition intervals reflected the frequency of read/SNP combination occurrences, which became sparser in regions of poorly assembled sequences in the D2 assembly. The result is a skewed distribution of sizes. The median size of transition intervals was ~6 kb, the mean was around ~150kb and the total transition interval space across the genome was found to be 362 Mb.

For our *de novo* genetic map marker positions were selected near the end of each transition interval. In this way, a marker represents the genotype at its location through to the next marker, with a small region of uncertainty at the end in the transition interval. Effectively each marker represents a unique genomic interval within the mapping population. In most cases, SNPs were genotyped multiple times within overlapping alignment reads. Care was taken to avoid selecting regions of rapid genotype switching due to residual heterozygosity or experimental noise. Any regions of residual heterozygosity were recognized by genotype variation among reads overlapping the same SNP and by rapid genotype switching among sequential SNPs. Rates for rapid genotype switching due to experimental noise, identified using the B6 parental strain samples, were low and avoided.

The transition intervals across all the selected BXD lines were combined to create a genome-wide map of its haplotype blocks. Using these *de novo* maps resolved most conflicts between genotype and phenotype (Supplemental Fig. S1). To estimate the functional extent of disagreement between genetic maps we characterized all H3K4me3 peaks with LOD > 25 (n = 807, mapped using the *de novo* genotypes). This class of H3K4me3 peaks offer the clearest test cases that allow strong separation in H3K4me3 level between B6 and D2 genotypes, driven by local variants. Of these *cis*-QTL, 69% (n = 560) had at least one strain with a disagreement between the local genotypes between genetic maps, after excluding heterozygous regions. Upon visual inspection, some genotyping differences were due to one marker positioned 5’ or 3’ to our called *de novo* markers, likely a result of improved resolution of our genetic maps. To exclude these cases, we next required that two markers in both directions from the QTL interval disagree between genetic maps, resulting in a broader region of discordance between maps. This resulted in 38% (n = 310, representing 198 unique genetic intervals) of *cis*-QTL with LOD > 25 having at least one strain with disagreement between genotypes. In all cases examined, the *de novo* genetic map led to better agreement between H3K4me3 level and genotype at *cis* QTL with LOD scores > 25 (Supplemental Fig. S1). Two strains, BXD81 and BXD100 consistently had large regions in disagreement between the two genetic maps. Maps from all RI lines used in this study were formatted for compatibility with R/qtl2 package and are available as Supplemental Table S7.

### Statistical analysis and QTL mapping

All statistical analyses were performed using R (http://www.R-project.org/). Differential H3K4me3 levels between parental strains for Figure 1, and for the *trans*-regulation test with BXD strains, were calculated using the exact test from the edgeR package (Robinson et al. 2010). All samples were normalized using the TMM method. P-values were adjusted using the false discovery rate method of Benjamini-Hochberg.

QTL analysis was performed using R/qtl2 package (Broman et al. 2003). Prior to mapping the matrix of samples-by-H3K4me3 level was TMM normalized and log_2_ transformed. Data exploration using hierarchical clustering found that replicate samples did not always group together (Supplemental Fig. S5A). This was found to be due to large variation in the H3K4me3 signal found at recombination hotspots (Supplemental Fig. S5B), suggesting that genetic replicates of juvenile mice coming from different litters might be at slightly different stages of meiosis. Although each litter of mice was collected at 14 dpp, there might be small variation in timing of meiotic entry due to genetic or environment (such as litter size, time of day of birth, or mouse room). Principle component analysis found that the first PC described the variation in H3K4me3 level at hotspots and hierarchical clustering after subtracting PC1 found that replicates of the same strain clustered together (Supplemental Fig. S5C). Furthermore, genome scans using PC1 as a phenotype did not identify and significant or suggestive QTL, further suggesting that timing of recombination-specific H3K4me3 is not a genetic trait.

Consequently, QTL mapping was performed on PC1-subtracted data. Single-QTL scans were performed with the scan1 function of R/qtl2 using a linear mixed model and accounting for kinship. We used a permutation approach to generate genome-wide FDR thresholds. 1000 permutations were performed by randomly permuting the samples labels, and an empirical maximum LOD score distribution was computed for each H3K4me3 peak. An empirical p-value was calculated for each peak by comparing the original LOD score for each peak with the empirical LOD score distribution, and FDR was computed across the whole dataset using the R package fdrtool (Strimmer 2008). For each H3K4me3 peak, only the QTL location with the highest LOD score was kept for further analysis (Supplemental Table S3).

### Network Analysis

For network analysis QTL scans were performed as above assuming an additive model and conditioning on the closest marker to the peak. Samples were permuted 2000 times for each H3K4me3 peak retaining LOD scores greater than the 95^th^ percentile of the empirical maximum. Peaks found to be in linkage disequilibrium with their *trans* marker, or FDR > 0.05, were filtered prior to network analysis, resulting in 2,689 *trans*-associated peaks. These were clustered into modules using a weighted gene co-expression network analysis (WGCNA) (Langfelder and Horvath 2008). To obtain tightly correlated modules, we used iterativeWGCNA (Greenfest-Allen et al. 2017), with the following settings: power = 12; minKMEtoStay = 0.7; minCoreKME = 0.7, and the remaining parameters set at default values. This resulted in 1,224 H3K4me3 peaks clustered into 14 tightly correlated modules (Supplemental Table S4). The first principal component of each module (termed eigengenes) is a representation of the summary co-expression pattern, and treated as module phenotypes for module-QTL analysis using R/qtl2.

### Annotation of H3K4me3 sites

Recombination hotspot locations in both B6 and D2 mice were established by mapping the locations of the DMC1-associated ssDNA that arises at meiotically programmed DNA double-strand breaks (DSBs) (Smagulova et al. 2011). We classified remaining sites by whether they corresponded to promoters, based on overlapping TSS, or to enhancers and CTCF sites using mouse ENCODE testis datasets (Shen et al. 2012), or to transposable elements.

### Motif Enrichment Analysis

Analysis of Motif Enrichment (AME in MEME Suite 4.12.0 (Bailey et al. 2009)) was used to identify motifs that are significantly enriched in module peaks. For controls, 10,000 sequences were randomly selected out of the total 67,110 H3K4me3 peaks. HOCOMOCO Mouse (v11 CORE) database was used to test for enrichment (Kulakovskiy et al. 2018). Motifs and their corresponding transcription factors, with adjusted p-value < 0.01 (Bonferroni), are reported in Supplemental Table S5.

### Measuring DMC1-associated single-stranded DNA fragments for DSB estimates

DMC1 ChIP was performed with isolated and crosslinked spermatocytes from B6 and D2 mice using an established method (Khil et al. 2012). Briefly, testes from adult B6 or D2 mice were crosslinked in 1% paraformaldehyde for 10 min, and then homogenized. Spermatocytes were washed with PBS and lysis buffers. Then, the chromatin was sheared to 500-1000 bp by sonication and incubated with DMC1 antibody (Santa Cruz, sc-8973) overnight at 4°C followed by a 2 hours incubation with Protein G beads (Life Technologies, 10004D). The beads were washed and the chromatin was eluted and reverse-crosslinked with 1% SDS, 0.1 M NaHCO3, 0.2 M NaCl at 65°C overnight. The libraries were constructed with TruSeq Nano DNA LD Library Prep Kit (Illumina, FC-121-4001, set A) using enzymes from New England Biolabs. DNA samples were sequenced on an Illumina^®^ platform, with 75 bp paired-end reads.

Detection of DMC1 Chip-Seq peaks across both B6 and D2 was accomplished by aligning two replicate paired-end B6 samples and two replicate paired-end D2 samples to their associated genomes, GRCM Build 38 (mm10) and the D2 *de novo* REL-1504-Assembly respectively, utilizing bwa version 0.5.10-tpx (Li and Durbin 2009). The fastq files were trimmed with Trimmomatic version 0.32 (Bolger et al. 2014) and any adapter sequences removed.

During sample preparation, DNA single strand fragments bound to DMC1 were enriched from background DNA by fast annealing short sequences of intra-molecular micro homology, followed by exonuclease removal of 3' overhang ends and polymerase fill-in of remaining 5’ overhang sequence (Khil et al. 2012). This generates 3’ and 5’ end homologous sequences that arise from the region of micro homology combined with filling in the 5’ overhangs. Sequencing reads were selected for the presence of these regions of 3’ and 5’ homology, and the homologous portions deleted. The trimmed reads selected in this way were then aligned to their corresponding parental genome assemblies. Because the majority of the insert sequences are shorter than the read lengths, the reads from one strand often contain a fill in sequence. The single stranded DNA reads without fill-in were then selected by evaluation of the alignment flags and retained. This created a new single-end fastq file that was subsequently aligned for peak calling. Peaks were called using MACS version 2.1 (Zhang et al. 2008) with the extsize option set to 800 in order to align reads into detectable clusters. Peaks were linked between B6 and D2 similar to methods applied to H3K4me3 reads as described above.

